# From Lab to Concert Hall: Effects of Live Performance on Neural Entrainment and Engagement

**DOI:** 10.1101/2025.04.03.646931

**Authors:** Arun Asthagiri, Psyche Loui

**Author notes:** Corresponding Author: Psyche Loui.

## Abstract

Live music performances continue to captivate audiences despite widespread availability of high-quality recordings, yet the neural mechanisms underlying this enhanced experience remain poorly understood. This study investigates the effect of live versus recorded music on neural entrainment using phase-based approaches. 21 participants listened to 2 live and 2 recorded performances of fast and slow movements of J.S. Bach’s works for the solo violin in a concert hall setting, while their EEG data were collected. Participants made behavioral ratings of engagement, spontaneity, pleasure, investment, focus, and distraction after each trial. Live performances were rated as more engaging, pleasurable, and spontaneous than recorded performances. Live trials showed significantly higher acoustic-EEG phase-locking than recorded trials in frequencies specific to the tempo of the excerpts. Furthermore, the effect of liveness on phase-locking was linked to increases in pleasure and engagement for live over recorded trials. Control analyses confirmed that the effects of liveness on phase-locking were not explained by low-level acoustic differences between performances. Altogether, results provide the first evidence that live music enhances cerebro-acoustic phase-locking, and that this enhanced entrainment underlies the heightened affective experience of live performance, supporting theories of music as a vehicle for social bonding through shared neural dynamics.

## Introduction

Music is often cited as one of life’s greatest pleasures, and despite the proliferation of audio and video streaming services, attending live music performances remains one of the most popular activities across age groups and cultures. A growing body of literature shows that liveness (i.e. real-time, in-person experiences) plays an important role in social interaction, mental wellbeing, and collaborative communication (Balters et al., 2023; Stieger et al., 2023). Audiences cite a variety of psychological reasons for being drawn to live performances, including social engagement and novelty seeking (Brown & Knox, 2017), though less is known about the neural dynamics underlying the perception of live music.

During live concerts, audiences synchronize at neural and physiological levels (Chabin et al., 2022; Czepiel et al., 2021; Tschacher et al., 2023), mirroring social synchrony found in nature (Greenfield et al., 2021; Strogatz, 2015). Synchrony through rhythmic movement in particular is linked to prosociality and social bonding (Cirelli et al., 2014; Fujiwara et al., 2020). This rhythmic synchrony can emerge from entrainment to musical rhythm (i.e., in dance, Bigand et al., 2024) because rhythm facilitates joint-action (Rabinowitch, 2023) and a pleasurable desire to move (Janata et al., 2012; Witek et al., 2014). While the impact of rhythm on synchrony and prosociality is taken as evidence that live music developed as a means for social bonding (Grahn et al., 2021; Savage et al., 2021), it raises the question of whether liveness itself impacts neural entrainment to rhythm.

In this vein, manipulations of social and affective contexts demonstrate that rhythms perceived as human-created elicit stronger effects of entrainment and liking relative to rhythms believed to be generated by non-human counterparts. Rhythmic entrainment is stronger when listening to music relative to a metronome (Stupacher et al., 2017). Behavioral research shows that rhythmic synchrony predicts likeability for human and not machine partners (Launay et al., 2014). Electroencepholography (EEG) studies show that neural entrainment correlates with the pleasurable urge to move only when melodies are human-performed (Cameron et al., 2019). By extension, we hypothesized that listening to live relative to recorded music would increase neural entrainment to rhythm and perceived pleasure and engagement.

Neural entrainment is the process through which neural oscillations align in phase and frequency with rhythmic input. Neural entrainment arises when periodic events reset the phase of neural oscillations and modulate the timescale of neuronal excitation (Obleser & Kayser, 2019). The resulting phase-alignment of neural oscillations with periodic auditory activity is thought to facilitate encoding and prediction of acoustic events (Large & Jones, 1999). Studies in selective attention have shown that directing attention toward an auditory stream strengthens neural-acoustic phase-alignment (Lakatos et al., 2008; Schroeder & Lakatos, 2009) and biases perception (Zion Golumbic et al., 2013). Thus, neural entrainment is considered to be a mechanism for auditory sensing that is guided by higher-level cognitive processes (Lakatos et al., 2019; VanRullen, 2016).

Identifying genuine neural entrainment poses methodological challenges (Duecker et al., 2024), for example in distinguishing sustained entrainment of neuronal oscillations from repeated evoked potentials (Novembre & Iannetti, 2018). Here we use neural-acoustic phase-locking (Lachaux et al., 1999) as a measurement of neural entrainment in the broad sense (Obleser & Kayser, 2019). Phase locking captures the cyclic alignment between neural and acoustic data at specific frequencies. Empirical and modeling studies suggest that neural–acoustic phase alignment represents neural entrainment to naturalistic speech and music (Doelling et al., 2019; Zoefel et al., 2018).

Features of speech and music are known to drive neural entrainment. In speech, quasi-rhythmic delta-theta modulations drive neural–acoustic phase-locking (Doelling et al., 2014; Luo & Poeppel, 2007; Peelle et al., 2013). In music, neural–acoustic phase-locking occurs at multiples of the beat rate (Tichko et al., 2022a), and depends on acoustic features like spectral complexity (Wollman et al., 2020) and individual differences in musical experience (Doelling & Poeppel, 2015; Harding et al., 2019). Neural activity at beat-related frequencies in music are predictive of the pleasurable urge to move or “groove” (Zalta et al., 2024). In sum, neural entrainment reveals a link between low-level auditory perception (Henry & Obleser, 2012), attentional modulation (Calderone et al., 2014), and sensorimotor engagement (Rosso et al., 2023), which makes it well-suited for understanding how the experience of live music engages the listening brain.

In the present study, we test whether the experience of liveness (i.e., the quality of an in-person performance, versus a recording by the same performer) influences neural entrainment to the acoustic signal for single listeners. We conceptualize the experience of liveness as the interplay between top-down effects from engagement with (awareness of, and attention towards) a live performer, and bottom-up effects from acoustic differences in the performance itself. Performance studies have shown that live relative to recorded concerts increase physical engagement (Swarbrick et al., 2019) and physiological synchrony (Kragness et al., 2023). Recent research has also shown that audience-performer interactions increase activity in limbic networks and strengthen brain-music correlations in a neurofeedback paradigm (Trost et al., 2024). However, it is not known whether the mere context and experience of the live performance alters neural entrainment to rhythm. In this <u>preregistered study</u>, we use EEG to test whether and how the presence and experience of live performances relative to recorded controls affect neural– acoustic phase-locking in a concert hall. We hypothesized that live over recorded music would (1) result in more positive reports of pleasure and engagement, (2) influence neural oscillations during listening, and (3) increase the strength of neural–acoustic phase-locking to rhythms in the music.

## Methods

### Participants

21 participants (11 female, mean age 22.9 yrs, range 18-31 yrs) were recruited from online surveys and advertisements from the New England Conservatory of Music community. All participants had a history of formal musical training (n=5, 3-9 yrs; n=16, > 10 yrs) in violin, other strings, winds or voice (n=7, 8, 3, 3 respectively) and at least two years of music theory training. Participants gave informed consent as outlined in IRB #18-12-13, and were compensated $25.

### Stimuli

Four excerpts were selected from J.S. Bach’s Solo Violin Sonatas and Partitas as stimuli for the experiment. Two fast movements (approx. 125 beats per minute) and two slow movements (approx. 50 bpm) were selected to match for style and structure within tempo conditions: Sonata No. 1 mov. 1—Adagio (slow), Sonata No. 2 mov. 1—Grave (slow), Partita No. 1 mov. 1— Preludio (fast), and Sonata No. 3 mov. 4—Allegro (fast). Two of these movements, one fast and one slow, were presented in live performance by the renowned violinist Joshua Brown. The other two movements were presented with a dual-speaker system (see Figure 1A). The order of live and recorded stimuli was alternated within each participant and counterbalanced across participants. The order of tempo was pseudorandomized such that every participant experienced all four Bach excerpts once each, and every participant experienced 4 conditions in a 2 x 2 design: Live-Fast, Live-Slow, Recorded-Fast, and Recorded-Slow.

**Figure 1.**
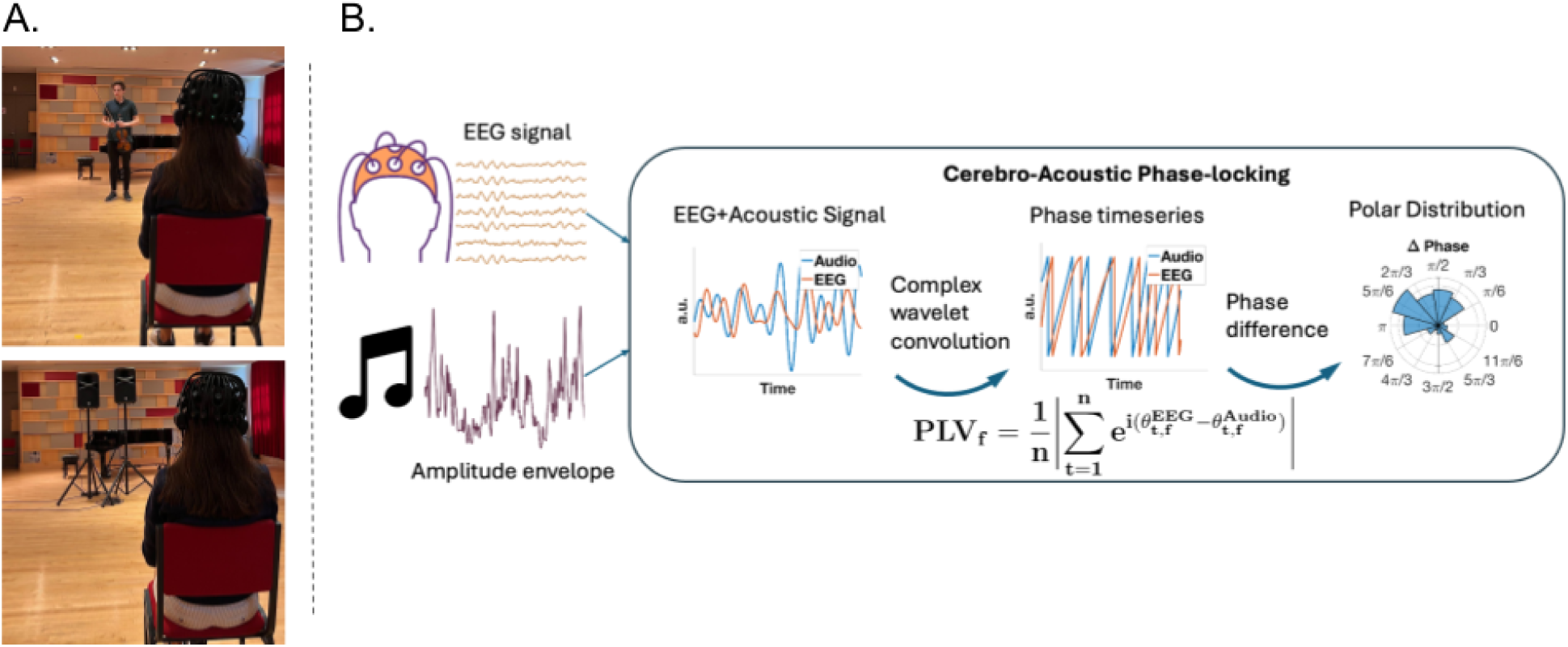
Data Collection and Analysis Methods. (A) Live and recorded performances were presented separately to 21 participants in a concert hall (eyes closed). EEG was recorded using a mobile system. Live and recorded trials were matched for loudness and acoustic source location. (B) Phase-locking and the temporal response functions revealed naturalistic Neural–acoustic phase-locking provided a measure of the phase-angle coherence between acoustic and EEG signals at a given frequency. **(Figure 1. Alt. text**

### Procedure

Participants listened to four excerpts (live/recorded x fast/slow) presented in a concert hall (Pierce Hall) at the New England Conservatory, while EEG was recorded with eyes closed. Two recorded performances were given by concert violinist and soloist, Joshua Brown, who played on a 1679 Guarneri Del Gesu violin. Two recorded excerpts were played from recordings made by the same violinist in the same concert hall prior to data collection.

After informed consent procedures, participants completed questionnaires on demographic information and standardized assessments of musical reward sensitivity (Extended Barcelona Music Reward Questionnaire, eBMRQ, Mas-Herrero et al., 2013; musical sophistication and engagement (the Goldsmiths Music Sophistication Index, Gold-MSI, Müllensiefen et al., 2014), The Synesthesia Battery (Eagleman et al, 2007), and the Adult ADHD Self-Report Scale (ASRS, Kessler et al, 2006). The experiment generally lasted 1 hour for each participant including EEG setup, questionnaires, and listening. Participants rated their listening experience of each excerpt (∼ 4 minutes) directly after presentation, reporting engagement, enjoyment, familiarity, pleasure, focus, investment, distraction (reverse-scored) and spontaneity.

### Data Acquisition

EEG data were recorded using a dry Quick-32r headset at 500 Hz and collected with the CGX custom acquisition software. Electrode impedances were kept under 300 kOhms. A CGX Wireless Stim Trigger device was used to align audio recordings with EEG data at the start of each trial.

All live and recorded performances were recorded for analysis using Pierce Hall’s DPA 4011C stereo microphone system. For two excerpts, backup recordings from a separate Blue Yeti microphone were used due to errors with the recording system.

A Yamaha 600S dual-speaker system was used for recorded excerpt playback for the first 11 participants, and a JBL Eon 500 was used as a comparable large dual-speaker PA system for the remaining 10 participants. The location of the playback system and performer were defined relative to the participant as 14 feet from the audio source (measuring speaker to ear). The bottom of the speakers was situated 4.5 ft above the ground to match the height of the violin position during live trials. The dual speakers were placed close together (approx. 1 ft 10 inches apart) to simulate the single sound source of the violin. Levels were matched in decibels between speaker and violinist prior to data collection by recording the violinist and the speaker separately and iteratively matching speaker levels, and adjusted per trial to minimize differences in amplitude between live and recorded performances.

### Analysis Plan

#### Behavioral data analysis

Participants gave 5-point Likert ratings after each musical excerpt on pleasure, enjoyment, engagement, familiarity, focus, investment, spontaneity, and distraction (distraction was reverse-scored). Principal component analysis (PCA) was conducted to find shared sources of variance across questionnaire items using the princomp function in R. The first principal component had strongest contributions from enjoyment and engagement, and weakest contributions from distraction and familiarity (S1). Projections of participant ratings onto the first principal component were thus used as a “pleasure-engagement” score for subsequent analyses. Pleasure-engagement scores were entered into a generalized linear mixed effects model (glmmTMB in R) to test for fixed-effects of tempo and liveness using participant as a random intercept.

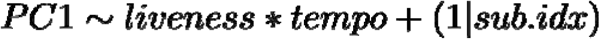

#### Audio preprocessing

Recordings from each trial were normalized for loudness using Matlab’s Audio Toolbox (v. 24.1) to obtain a final gain of −20 LUFS. Stereo recordings were converted into mono tracks by averaging across channels for subsequent analyses.

The following acoustic preprocessing steps are consistent with the literature and performed to simulate the filtering of the cochlea (Harding et al., 2019; Vanden Bosch Der Nederlanden et al., 2020a; Wollman et al., 2020). These steps were carried out with MIR Toolbox using default parameters unless otherwise noted (v. 1.8.2, Lartillot et al., 2008): Normalized audio was filtered across 50 logarithmically-spaced frequencies using a gammatone filterbank (mirfilterbank). The amplitude envelope was computed within each frequency band using a lowpass FIR filter (mirenvelope), and summed across frequency bands (mirsum). The resulting acoustic signal was downsampled to 500 Hz to match the EEG sampling rate for phase-locking analysis. Peaks in the frequency spectrum of the amplitude envelope reflected rhythmic activity in slow and fast excerpts (S2).

#### EEG preprocessing

EEG data was trimmed to the onset of each excerpt using acoustic triggers presented at the beginning of each excerpt. Data was preprocessed using a custom MATLAB script that called EEGLab functions (Delorme & Makeig, 2004). Channel locations were defined according to the international 10-05 standard with the MNI-BEM template. Data was referenced to the right earlobe (A2) and filtered with a high-cutoff of 58 Hz and a low-cutoff of 0.2 Hz. The low-cutoff was chosen to include the beat rate of the slow excerpts (∼50 bpm or .8 Hz). Noisy channels were identified and rejected when the signal was correlated at r<0.5 from the surrounding channels’ random sample consensus (RANSAC) estimate; had a flatline duration of more than 0.5 seconds; or had noise which reached more than 4 standard deviations of the signal. Artifact Subspace Reconstruction (ASR) was used to extrapolate data epochs where the variance was larger than 10 standard deviations from the calibration data (Chang et al., 2018; Tichko et al., 2022a). After visual inspection of the data to remove additional outlier channels, all rejected channels were interpolated spherically. Independent Component Analysis was computed on the EEG data and components that were explained more by non-brain sources than within-brain sources were identified and removed using EEGLab’s ICLabel method.

#### Neural–Acoustic Phase-Locking Analyses

Neural entrainment due to naturalistic music listening was operationalized using the phase-locking value (Figure 1B). In phase-locking analyses, the phase-coherence between the audio and brain stimulus is computed at specific frequencies, capturing continuous phase-adjustments of neural oscillations in response to acoustic cues. This reflects the theoretical assumption that neural responses to acoustic rhythms involves phase-frequency adjustments of neural oscillations (Obleser & Kayser, 2019).

The complex-valued signal was computed from both the real-valued cochlear-filtered audio and EEG signals using complex Morlet wavelet convolution. A wavelet of 5 cycles was selected, which is consistent with 3-7 cycles typically used in the neural–acoustic entrainment literature (Cohen, 2014; Doelling & Poeppel, 2015; Harding et al., 2019; Tichko et al., 2022b; Wollman et al., 2020).

Element-wise differences between phase-angle time series were calculated for linearly spaced frequencies from 0.2 to 20.2 Hz with a frequency increment of 0.2 Hz. The phase-locking value was obtained at each frequency of interest by computing the magnitude of the mean complex vector across samples (see equation below). A single phase-locking value was calculated per frequency, per electrode, and per trial.

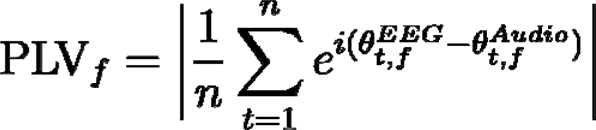

(Lachaux et al., 1999) Phase-locking values were averaged across the three frontocentral electrodes (F3, Fz, F4), which elicited highest coherence and aligned with the literature (Harding et al., 2019; Vanden Bosch Der Nederlanden et al., 2020; Wollman et al., 2020). Generalized linear mixed effects models (log link function) were separately applied to fast and slow conditions to model effects of live versus recorded performances on phase-locking strength over frequencies, accounting for excerpt as a fixed effect and participant as a random effect (model equation below). The model was run separately for each frequency, yielding p-values that were corrected for multiple comparisons using Benjamini & Hochberg FDR. Finally, we performed a confirmatory analysis using the same model on phase-locking values averaged across the significant contiguous frequency range for the fast excerpts.

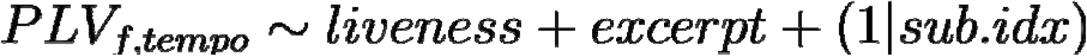

#### Brain-Behavior Relationships

To assess relationships between the strength of phase-locking and the perceived effect of liveness as operationalized by the behavioral ratings, we related the within-subject difference between the average phase-locking value over significant frequencies in the fast excerpts (the outcome variable in the confirmatory PLV analysis) to the within-subject difference in pleasure-engagement scores (the outcome variable in the behavioral analysis), using a generalized linear model. (No random effects were included at the participant level because the regressor and outcome variables were derived from within-participant differences.)

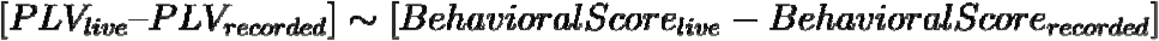

Panel A contains pictures showing the concert hall setup of the live condition and recorded conditions. The live condition picture has a violinist in the center frame and a participant with an EEG cap. The recorded condition picture has the dual speaker system in the center frame. Panel B shows the phase-locking analysis pipeline. The EEG and acoustic signals are transformed from a time-based representation to a phase-based representation using complex Morlet wavelet convolution. Phase-locking is derived from the phase difference between acoustic and EEG signals at a given frequency.)

#### Acoustic Control Analyses

Using MIRToolbox (Lartillot et al., 2008), we extracted six acoustic features from the recordings of each trial (using the normalized recorded audio after Audio preprocessing, see above). Of the six features, three features characterized changes in the time domain, i.e. “temporal features”: RMS loudness (mirrms), pulse clarity (mirpulseclarity), and spectral flux (mirflux), and three features characterized timbral components, i.e. “spectral features”: spectral centroid (mircentroid), brightness (mirbrightness), and zero-crossings (mirzerocrossings). Each of the six acoustic features was obtained for each audio recording, resulting in 504 total musical feature data points (21 participants x 4 performances each x 6 features per recording). We used Principal Component Analysis (princomp in R) to reduce the dimensionality of these acoustic descriptors, and to identify acoustic commonalities and differences across live and recorded performances of the same excerpts. Factor rotation was performed on the principal component scores using the varimax function in R to maximally orthogonalize loadings and potentially highlight differences between tempo and liveness conditions.

After determining that the first principal component of the acoustics differentiated live versus recorded performances, we tested whether this component significantly improved the fit of the generalized mixed effects models that were previously used. We performed likelihood ratio tests using the lmtest package in R. We compared reduced models from the aforementioned behavioral and neural analyses against full models that additionally included the first acoustic principal component scores.

## Results

### Pleasure and Engagement are Higher during Live Music Listening

Our first hypothesis was that live music would elicit more positive behavioral ratings than recorded music overall. Indeed, participants rated live trials higher than recorded trials in engagement, enjoyment, pleasure, focus, investment and spontaneity (Figure 2A). Ratings of familiarity and distraction were similar between live and recorded performances. A principal component analysis (PCA) was conducted to examine shared sources of variance in survey responses (see Behavioral data analysis). This PCA revealed two principal components, defined hereafter as “pleasure-engagement” (PC1) and “distraction-familiarity” (PC2) (Figure S1 reports loadings for each questionnaire item). A generalized linear mixed effects model was used to assess effects of liveness and tempo on both PC scores. Live music elicited significantly higher levels of pleasure-engagement than recorded music (β*=0.25, SE=0.046, z=5.51, p<.001*, Figure 2B) with a non-significant tempo interaction (β*=-0.039, SE=0.065, z=-0.60, p=. 55*), suggesting that regardless of tempo, live music elicited a more positive overall experience than recorded music.

**Figure 2.**
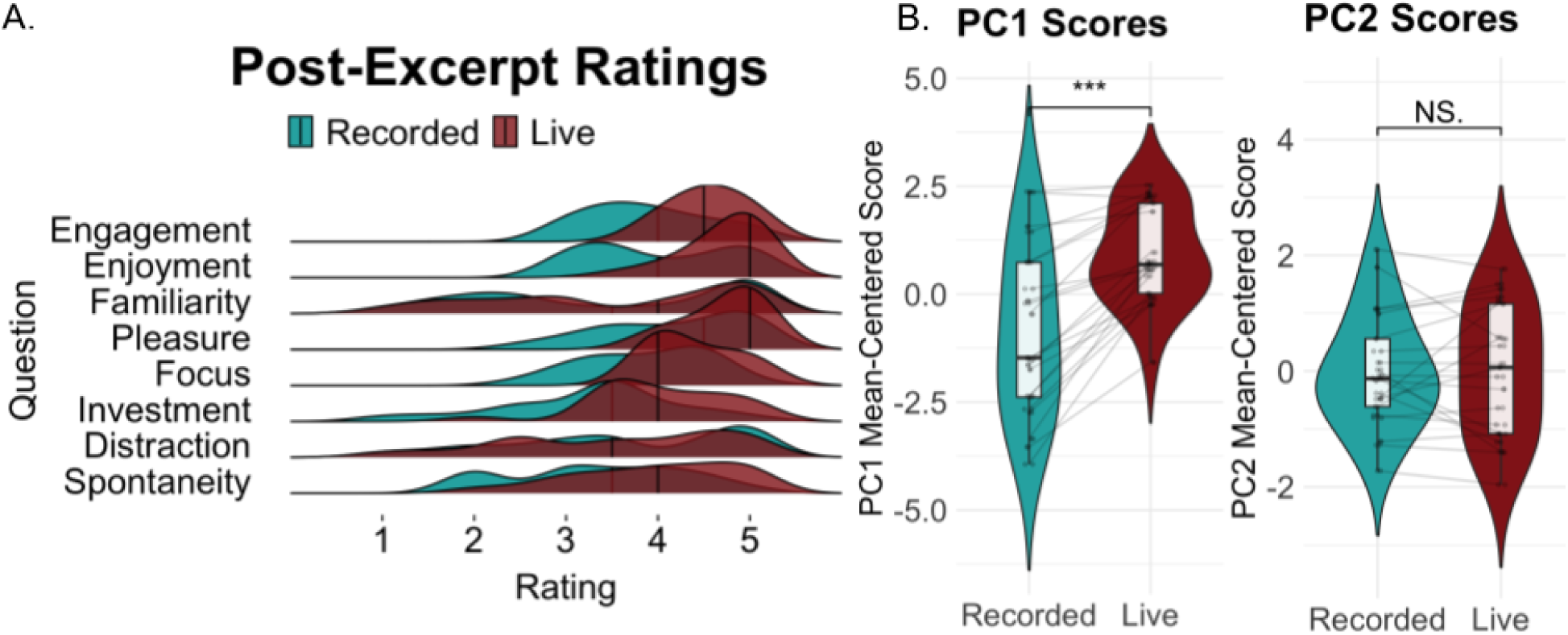
Behavioral responses to post-excerpt questionnaires. (A) Live compared to recorded trials were rated higher on engagement, enjoyment, pleasure, investment, and spontaneity, and similarly for familiarity and distraction. (B) The first principal component, which summarized a single “pleasure-engagement” dimension, was significantly higher for live over recorded performances (F(1,80)=12.72, p<.001). The second principal component summarized a “distraction-familiarity” dimension and did not differ significantly between live and recorded performances. **(Figure 2. Alt. text**

Panel A is a ridge plot of behavioral ratings for the 8 questionnaire items (engagement, enjoyment, familiarity, pleasure, focus, investment, distraction, and spontaneity) grouped by live and recorded conditions. Panel B shows results from the principal component analysis of behavioral ratings in a violin plot comparing PC1 scores between live and recorded conditions. Panel C shows a violin plot comparing PC2 scores between live and recorded conditions.)

### Liveness is Linked to Stronger Phase-Locking at Salient Acoustic Frequencies

Line plots showing neural–acoustic phase-locking values plotted over frequencies. Frequencies with significantly different phase-locking between live and recorded performance are marked by vertical red lines and shaded regions. Topographic plots are shown next to significant frequencies and depict differences in phase-locking between live and recorded performance over electrodes. Panel A shows phase-locking values for the slow excerpts, separately for live and recorded conditions. A vertical red line is marked at 3.4 Hz. Panel B shows phase-locking values for the fast excerpts, separately for live and recorded conditions. There is a shaded area between 7.6 Hz and 8.4 Hz.)

Phase-locking was used as a measure of audio-brain coherence across a range of frequencies (see phase-locking methods). Figure 3A shows phase-locking in slow-tempo trials, and Figure 3B shows phase-locking in fast-tempo trials. Peaks in phase-locking occurred in the delta and theta ranges for fast and slow excerpts, reflecting delta and theta rhythms in the acoustics themselves (S2). The delta range (.5-4 Hz) included the beat rate and note rate of the slow excerpts (beat rate: 50 bpm = .83 Hz; note rate: 4*50 bpm = 3.33 Hz), whereas the upper-theta range included the note rate of the fast excerpts (note rate: 4*125 bpm = ∼8.3 Hz).

**Figure 3.**
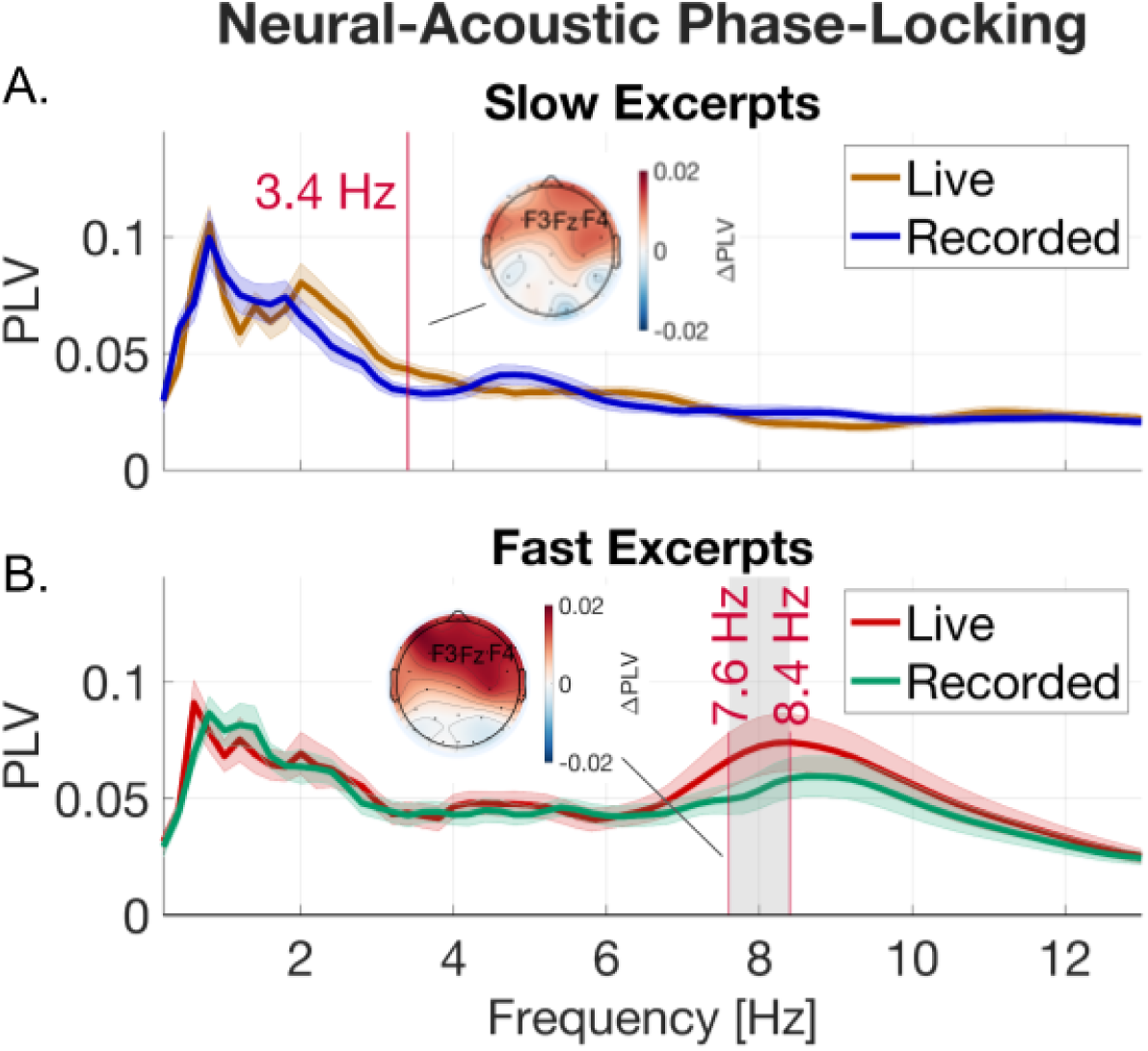
Cerebro-acoustic phase-locking differs across tempo and liveness. Phase-locking values between neural and acoustic activity plotted across a range of frequencies for (A) slow and (B) fast excerpts (shaded regions are +-1 standard error). Frequencies demarcated in red contained significant differences in phase-locking between live and recorded performances (generalized mixed effects models; FDR corrected p<.05). Topo plots show average differences in phase-locking strength over significant frequencies for live over recorded performances. **(Figure 3. Alt. text**

We tested for differences between live and recorded conditions using generalized linear mixed effects models at each frequency and corrected for multiple comparisons across frequencies using FDR. The model included liveness and excerpt as fixed effects and participant as a random effect (see Methods). Live conditions showed significantly stronger phase-locking than recorded conditions in the delta range (3.4 Hz) for the slow excerpts and in the upper-theta frequency range (7.6-8.4 Hz) for the fast excerpts (*p<.05, FDR corrected*). Frequencies with significantly higher phase-locking during live over recorded performance also corresponded to the note rates of the slow and fast excerpts (3.33 Hz and 8.3 Hz respectively).

We performed a confirmatory test on the average phase-locking value across this significant frequency range for the fast excerpts using the same model regressors. This revealed a significant effect of both excerpt and liveness on phase-locking strength (Liveness: β*=0.27, SE=0.08, z=3.36, p<.001;* Excerpt: β*=0.43, SE=0.08, z=5.42, p<.001*). Subsequent brain–behavior analyses focused on phase-locking over this upper-theta frequency range (7.6-8.4 Hz) of the fast excerpts as it showed a robust peak in phase-locking that aligned with the note-rate of these excerpts and significantly differed between live and recorded conditions.

Scatter plot showing participant-level differences between phase-locking during live and recorded fast excerpts on the x-axis and pleasure-engagement scores for the same excerpts on the y-axis. Histograms of the phase-locking differences and pleasure-engagement differences are shown next to the x and y axes respectively. Dashed vertical and horizontal lines mark the axes themselves (and represent the 0 values for both variables). A regression line is plotted through the scatter points showing a positive linear relationship between variables.)

### The Effect of Liveness on Neural Entrainment Predicts Pleasure-Engagement

Given neural differences in phase-locking as well as behavioral differences in pleasure-engagement between live and recorded performances, we performed a follow-up analysis to relate within-participant phase-locking during music listening to behavioral ratings post-listening. First, we tested whether differences between live and recorded performances in the mean phase-locking value across the significant frequency range for the fast excerpts (7.6-8.4 Hz) predicted pleasure-engagement differences between the same live and recorded performances. Results from a generalized mixed effects model revealed that phase-locking differences were significantly associated with pleasure-engagement differences (β*=2.85, SE=.84, z=3.40, p<.001*, Figure 4), indicating that relative to recorded music, the strength of neural entrainment during live performances was significantly associated with the perceived engagement after listening.

**Figure 4.**
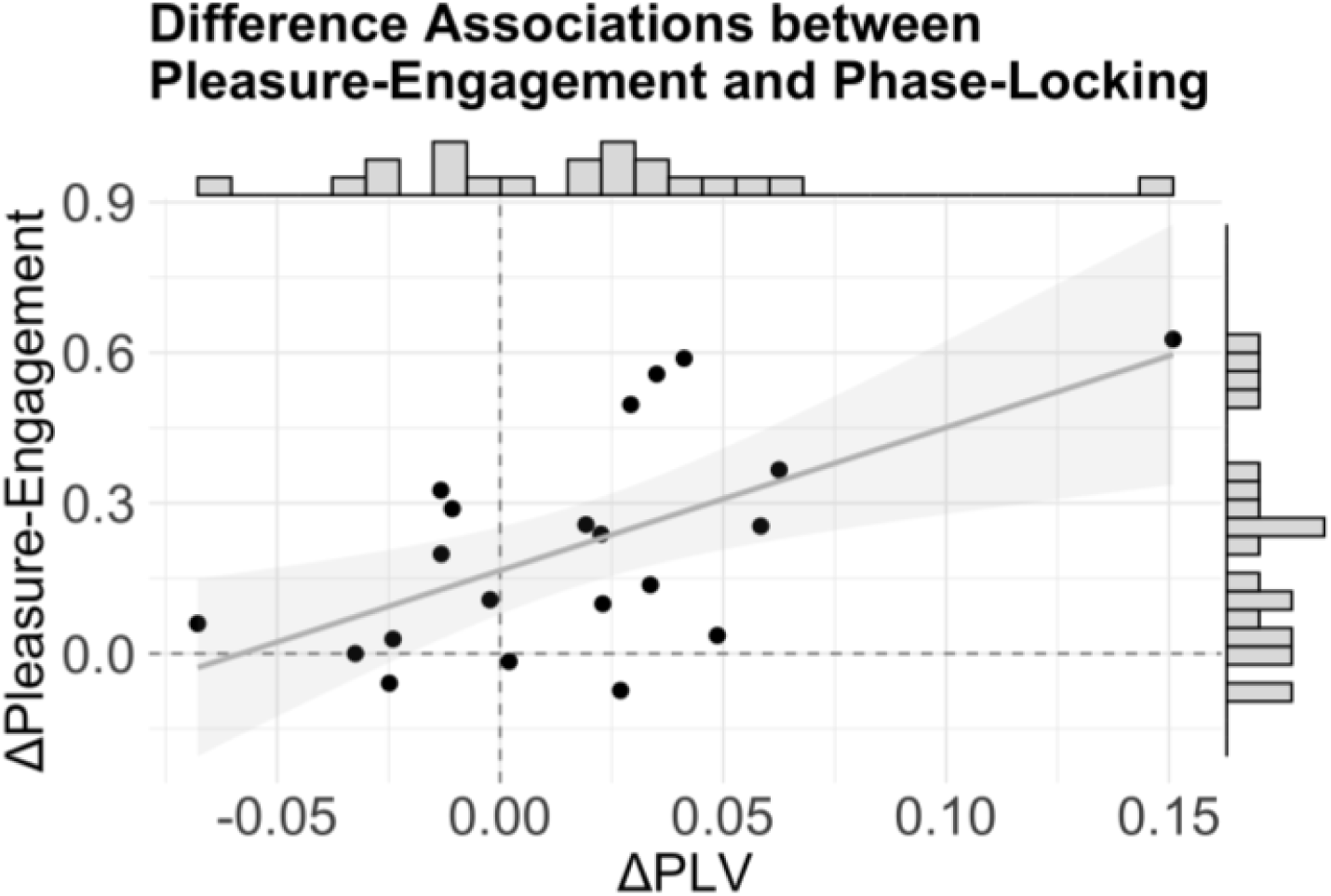
Phase-locking differences between live and recorded music predict differences in pleasure-engagement ratings. There was a significant relationship between the difference in phase-locking strength and the difference in pleasure-engagement ratings between live and recorded performances for fast excerpts (p<.001). Differences in pleasure-engagement (PC1) scores versus average phase-locking values between live and recorded performances are plotted. Points on the scatter plot denote single participants. Histograms show the marginal distributions of rating and PLV differences. **(Figure 4. Alt. text**

### Behavioral and Neural Effects of Liveness Extend Beyond Acoustic Differences

To what extent were the relationships between liveness, neural entrainment, and pleasure-engagement driven by differences in low-level acoustics as opposed to the participant’s top-down perception? While our study did not separately manipulate top-down perceptual bias and bottom-up sensory input, we did record all live and recorded performances and were thus able to perform follow-up analyses to test if any salient acoustic features may have driven differences between live and recorded performances. To quantify acoustic differences between live and recorded performances at the excerpt-level, we extracted low-level signal features from the normalized audio that were made available with MIRToolbox (Lartillot et al., 2008). These included both spectral features (spectral centroid, brightness, zero-crossings) and temporal features (RMS energy, pulse clarity, and spectral flux) (Figure 5A). Each of these features was summarized across an excerpt as a single value for each trial. These features were then inputted into a PCA to reveal shared sources of acoustic variance (Figure 5B).

**Figure 5.**
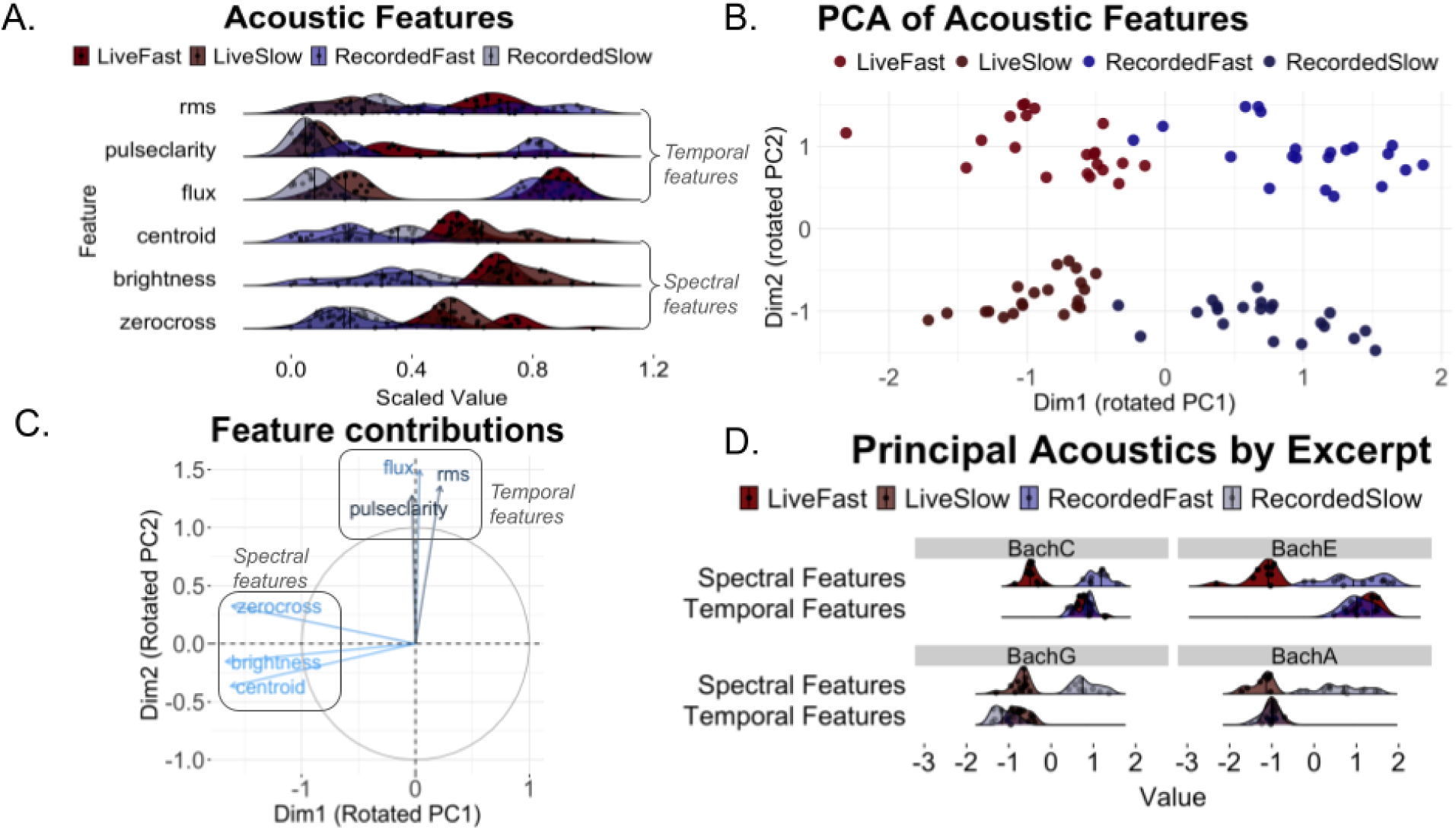
Acoustic Features of Live and Recorded Performance. (A) Density plots of acoustic features over trials show acoustic features that (B) distinguish liveness and tempo conditions after PCA. (C) Relative loadings reveal orthogonalized contributions of spectral (centroid, brightness, zero crossings) and temporal (RMS, spectral flux, pulse clarity) features.(D) Rotated PCs plotted within each excerpt highlight how spectral features differentiates liveness. **(Figure 5. Alt. text**

We found that the rotated PC1 and rotated PC2 had distinct contributions from spectral and temporal features respectively (Figure 5C). The spectral dimension (PC1) differentiated between liveness conditions whereas the temporal dimension (PC2) differentiated between tempo conditions (Figure 5B, 5D). The spectral dimension (PC1) was sensitive to timbral differences between the acoustic violin and the speaker, which were apparent to listeners even after controlling for overall amplitude (see methods). The variance in PC1 scores was not larger for live than for recorded trials (variance across live trials: 0.033; variance across recorded trials: 0.074), suggesting that timbral characteristics of live performances were consistent across trials.

Panel A is a ridge plot where rridges map onto distinct acoustic features (RMS, pulse clarity, spectral flux, spectral centroid,brightness, and zerocrossings). Each ridge shows distributions of acoustic feature values across trials. Separate distributions are shown for each of four conditions: Live Fast; Live Slow; Recorded Fast; Recorded Slow. Panel B is a scatter plot showing acoustic principal component scores for each trial across the spectral (PC1) and temporal (PC2) principal component dimensions. Points are colored by condition. Panel C is a circular plot showing loadings for each acoustic feature across principal component dimensions (x-axis is the rotated principal component 1; y-axis is the rotated principal component 2). Panel D shows ridge plots for spectral features (PC1) and temporal features (PC2) using separate panels for each excerpt (Bach C; Bach E; Bach G; Bach A). Individual ridge plots are grouped by condition.)

Lastly, we considered whether acoustic differences between live and recorded performances could have driven behavioral and neural outcomes in our previous models. We entered the first principal component of the acoustics, which differentiated between live and recorded performances as shown above, into the generalized mixed effects models used for the previous analyses. Likelihood ratio tests on reduced versus full generalized mixed effects models revealed no significant improvement in the model fit for pleasure-engagement scores (*df=1,* χ*2=1.42, p=0.23*) or for phase-locking values (*df=1,* χ*2=0.613, p=0.43*). Reduced model: tempo and liveness as regressors; full model: acoustics as an additional fixed effect regressor. Similarly, we added the first principal component of the acoustics into the phase-locking versus pleasure-engagement model, and found that PC1 acoustic differences between live and recorded performances did not significantly improve the model fit (*df=1,* χ*2=1.81, p=.18*). Together, these results suggest that timbral differences alone are insufficient to explain the effect of liveness on pleasure-engagement and neural entrainment. In light of these control analyses, our findings may predominantly be driven by top-down awareness of liveness (the top-down knowledge and awareness of a live performer) rather than bottom-up acoustic differences between live and recorded trials.

## Discussion

We provide a novel account of the effects of live music on neural entrainment. Musically-trained participants listened to live and recorded performances of slow and fast excerpts from the solo violin works of J.S. Bach presented in a concert hall by world-renowned violin soloist Joshua Brown. EEG was recorded during music listening, which was used to measure phase-locking between neural and acoustic signals. We demonstrated that (1) neural–acoustic phase-locking was guided by musical rhythmicity during naturalistic solo violin listening in a concert hall; (2) phase-locking was stronger during live over recorded performances; (3) pleasure and engagement differences between live and recorded performances covaried with phase-locking differences; (4) timbral differences between live and recorded mediums were not sufficient to explain behavioral or neural results. Together, results suggest that neural entrainment to musical rhythm increases during live over recorded music and accompanies pleasurable affect.

### Live Music Increases Phase-Locking to Musical Rhythm

We measured phase-locking between the acoustic envelope of live and recorded performances and the EEG activity of single listeners over a range of frequencies (.2-20.2 Hz). Neural–acoustic phase-locking was strongest at rhythmically salient frequencies in the delta and theta ranges for both live and recorded music, consistent with prior work showing phased-locked neural oscillations to speech and music across delta and theta frequencies (e.g., Peelle et al., 2013; Tichko et al., 2022b). For live over recorded music, phase-locking was stronger in the delta range for slow excerpts (3.4 Hz) and in the upper-theta range for fast excerpts (7.6-8.4 Hz), suggesting that the experience of liveness impacted the strength of acoustic and brain coherence.

The present study examines the experience of live music in a uniquely experimentally controlled manner in order to isolate the experience of liveness from social, visual, or acoustic variables. We minimized social effects between multiple performers or audience members by including only a single performer and a single audience member. We controlled for visual input by asking participants to listen with eyes closed. We also performed control analyses on salient acoustic features of live versus recorded performances to show that results were not driven by acoustic differences between the acoustic violin and the speaker system. Finally, recordings were made by the same performer that gave the live performances. Altogether, our approach suggests that liveness, even when experienced one-on-one, impacts neural–acoustic entrainment to musical rhythm.

Though our manipulation consisted of both acoustic and perceptual differences between live and recorded performances (see Figure 5), control analyses suggest that stronger neural–acoustic phase-locking occurs from live music not because of timbral differences, but because of the top-down awareness of the human performer. While previous studies report bottom-up effects of acoustics on neural–acoustic phase-locking e.g., melodic and spectral features (Vanden Bosch Der Nederlanden et al., 2020; Wollman et al., 2020), the top-down effect of liveness is novel and suggests a higher-level modulatory pathway for neural entrainment, akin to selective attention (Calderone et al., 2014). Future studies may trace the neural mechanisms that link liveness to neural entrainment by separately testing the roles of induced affect, attention, and social cognition during live music listening.

The impact of live music on neural entrainment has important implications. First, we extend findings from Trost et al., 2024, who focused on neural correlates of live performance feedback loops, showing that the effect of a live performer on acoustic–brain coherence does not rely on explicit neural feedback. Second, by recording acoustic data as well as neural activity with high temporal resolution, we extracted relationships between neural and acoustic activity that were tied to the frequencies of musical rhythms (i.e., delta and theta ranges). Since the relationship between rhythm in music and the pleasurable desire to move is well-established (e.g., Benson et al., 2024; Witek et al., 2014), our results provide a neural basis for how rhythms in music may drive audience members to exhibit more intense movements during live versus recorded concerts (Swarbrick et al., 2019). Third, increased phase-locking at rhythmic frequencies during live performance supports the notion that rhythmic entrainment is a cornerstone of live social bonding through music (Cirelli et al., 2018; Grahn et al., 2021; Savage et al., 2021). This is in line with theories of social resonance and collective effervescence in group dynamics which posit that neural synchrony yields behavioral synchrony, which is a mechanism for social understanding (Durkheim & Fields, 1995; Wheatley et al., 2012).

### Individual Differences in Phase-Locking Associate with Ratings of Pleasure and Engagement

While cerebro-acoustic entrainment has been established as a mechanism for auditory processing and attentional selection (Henry & Obleser, 2012; Lakatos et al., 2008), less is known about how entrainment interacts with the perceived pleasure of acoustic stimuli such as live music. Given that behavioral results revealed more positive affect for the live condition compared to the recorded condition, we looked at whether variability in the strength of neural entrainment explained differences in the subjective evaluation of live music. Related work has compared neural entrainment to human-performed versus mechanical rhythms, and found that pleasure and entrainment are correlated only in human-performed but not mechanical music (Cameron et al., 2019). In live settings, perceived pleasure is linked to the strength of inter-subject theta coherence in the audience (Chabin et al., 2022). Our results show that the difference in how listeners report pleasure and engagement between live and recorded performances is associated with their difference in neural–acoustic phase-locking for the same performances. This association between individual differences in brain and behavior—that is, between increased pleasure-engagement ratings and increased phase-locking for live over recorded performance— suggests that neural entrainment may be an indicator of affective experience in music.

Multiple neural pathways might explain how the top-down awareness of a live performer impacts both neural entrainment and pleasure. Attention is one such pathway. Attention is known to influence the fidelity of auditory encoding by strengthening neural entrainment to an acoustic stream (Calderone et al., 2014; Lakatos et al., 2019). During live music, the awareness of a live performer may increase attentional engagement toward the acoustic input. This increase in attention may provide top-down modulation on neural–acoustic phase-locking, resulting in better encoding of the musical stimulus. Thus, increased attentional engagement during live music may alter the pleasure of a listener by influencing neural entrainment and acoustic encoding. Conversely, pleasure itself may factor into the strength of neural entrainment. Musical pleasure is linked to connectivity between the auditory and reward systems (Belfi & Loui, 2020; Salimpoor et al., 2013). Arousal responses in listeners scale with specific sensitivity to musical rewards (Mas-Herrero, Zatorre, et al., 2014). Similarly, attention and engagement might arise from the specific pleasure associated with liveness. Sensitivity to the pleasure of live music might mediate the effect of liveness on neural entrainment. This would explain the relationship between individual differences in neural entrainment and pleasure between live and recorded performances. Treating neural entrainment as an interplay between bottom-up and top-down features of auditory and reward processing may reveal mechanistic insight into how music (and live music specifically) impacts neural entrainment.

### Conclusions

Together, the study provides novel insights into the effects of live music on neural entrainment in a controlled yet naturalistic concert hall setting. Employing phase-based methods for relating continuous EEG and acoustic signals, our results suggest that live over recorded music enhances neural–acoustic phase-locking to musical rhythms along with the pleasure and engagement of listeners. In the digital age of art consumption, the role of liveness is increasingly debated (Auslander, 2022). Our results reveal a neural correlate of liveness in the dynamics of music listening, providing the first evidence that liveness impacts neural entrainment to music. Future work may further uncover how the various social and affective dimensions of liveness manifest themselves in neural and physiological dynamics during live music listening.

## Data Availability

Code and data will be made available upon reasonable request to the corresponding author.

