## Supplementary Materials for "From Lab to Concert Hall: Effects of Live Performance on Neural Entrainment and Engagement"

### S1. PCA Loadings For Post-Excerpt Questionnaire Data

Participants gave ratings on engagement, enjoyment, familiarity, pleasure, focus, investment, and distraction, which were subjected to a principal components analysis (PCA). The first principal component scores were referred to as “pleasure-engagement” ratings. Loadings of the individual questionnaire items for this first principal component are shown below.
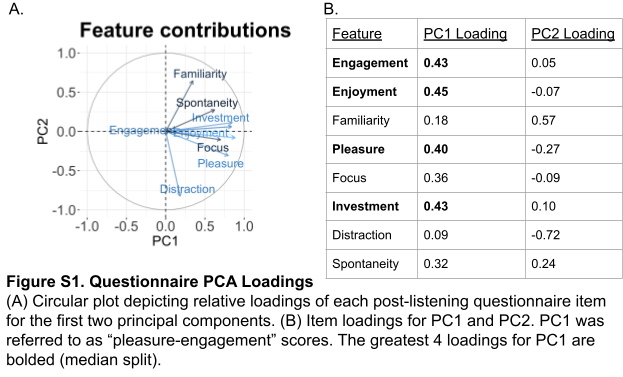

### S2. Acoustic Spectra of Slow and Fast Excerpts

Cochlear-filtered audio was passed through a fast Fourier transform (FFT) to obtain acoustic spectra for slow and fast excerpts. These spectra revealed rhythmic activity in the acoustics across delta and theta frequency ranges. The beat rate and note rates of the slow (~50 bpm) and fast (~125 bpm) excerpts, whose tempi were set prior to data collection, appear as peaks in the amplitude spectra: the beat rate of the slow excerpts (.83 Hz) appears as a peak in the low-delta range and the note rate of the fast excerpts (8.33 Hz) falls within a peak in the upper-theta range.

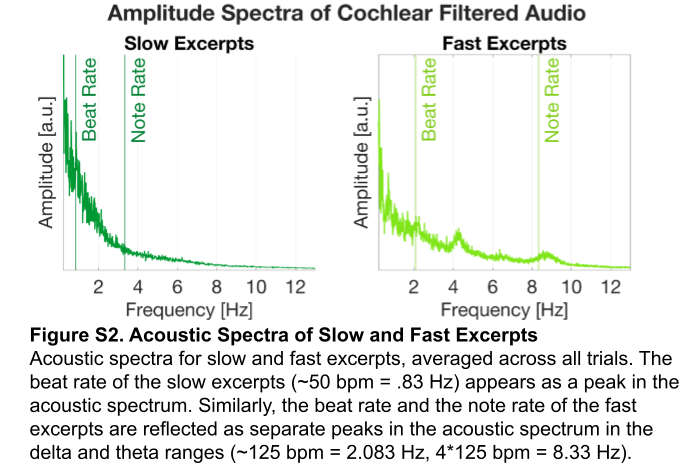
